# Inspiratory rhythmogenic activity is burst-independent and opioid-sensitive

**DOI:** 10.1101/665034

**Authors:** Xiaolu Sun, Carolina Thörn Pérez, D Nagaraj Halemani, Xuesi M. Shao, Morgan Greenwood, Sarah Heath, Jack L. Feldman, Kaiwen Kam

**Affiliations:** Department of Neurobiology, David Geffen School of Medicine at UCLA, Los Angeles, CA 90095, USA; Department of Cell Biology and Anatomy, Chicago Medical School, Rosalind Franklin University of Medicine and Science, North Chicago, IL 60064, USA; RFUMS/DePaul Research Internship Program, Rosalind Franklin University of Medicine and Science, North Chicago, IL 60064, USA

**Author notes:** Equal contribution (Xiaolu Sun, Carolina Thörn Pérez). Corresponding author and Lead Contact: Kaiwen Kam, Ph.D., Department of Cell Biology and Anatomy, Chicago Medical School, Rosalind Franklin University of Medicine and Science, 3333 Green Bay Rd., North Chicago, IL 60064-3095, USA.

**Keywords:** opioids, breathing, rhythm generation, central pattern generator

## Abstract

How mammalian neural circuits generate rhythmic activity in motor behaviors, such as breathing, walking, and chewing, remains elusive. For breathing, rhythm generation can be localized to a brainstem nucleus called the preBötzinger Complex (preBötC). Rhythmic preBötC population activity consists of small amplitude burstlets, which we hypothesize are rhythmogenic, and larger inspiratory bursts, which drive motoneuronal activity. If burstlets are rhythmogenic, opioids, analgesics that can cause profound respiratory depression, should similarly reduce burstlet frequency. In conditions where burstlets were separated from bursts in medullary slices from neonatal mice, the μ-opioid receptor (μOR) agonist DAMGO decreased burstlet frequency. DAMGO-mediated depression was abolished by genetic deletion of μORs in a glutamatergic preBötC subpopulation and was reduced by Substance P, but not blockade of inhibitory synaptic transmission. Our findings suggest that rhythmogenesis need not rely on strong bursts of activity associated with motor output and point to strategies for ameliorating opioid-induced depression of breathing.

## Introduction

Rhythmic motor behaviors in mammals, such as breathing, walking, and chewing, are controlled by neural circuits that determine the period of the movement and shape the pattern of motor activity. Despite the apparent simplicity of the computation, the mechanisms underlying rhythmogenesis in these circuits are unknown. Generation of breathing rhythm is localized to a compact nucleus in the ventrolateral medulla, the preBötzinger Complex (preBötC; Smith et al., 1991). In particular, two overlapping subpopulations of glutamatergic preBötC neurons, those expressing the neurokinin-1 receptor (NK1R) and those derived from progenitors that express the transcription factor Dbx1 (Dbx1^+^), are necessary for inspiratory rhythmogenesis (Bouvier et al., 2010; Gray et al., 2010; Gray et al., 2001).

Until recently, rhythmic bursts of preBötC population activity were considered unitary events that determined both the timing and pattern of periodic inspiratory movements. However, we recently showed that these preBötC bursts consist of two separable components: i) large amplitude inspiratory bursts that are transmitted via premotoneurons to inspiratory motoneurons to activate muscles that produce inspiratory airflow; and ii) low amplitude burstlets that appear as preinspiratory activity when preceding these bursts and are not normally seen in motor nerve or muscle activity (Kam et al., 2013a). Because preBötC rhythmic activity can consist solely of burstlets that do not produce motoneuronal output, we hypothesize that burstlets are rhythmogenic (Kam et al., 2013a). A strong prediction of this hypothesis is that manipulations that affect inspiratory burst frequency should have similar effects on preBötC burstlet frequency.

To test this prediction, we examined the effects of opioids, analgesics that depress breathing at high doses, on bursts and burstlets and tested opioidergic interactions with other neurotransmitters and neuromodulators that affect preBötC neuronal excitability. We bath-applied [D-Ala^2^,N-MePhe^4^,Gly-ol^5^]-enkephalin (DAMGO), a potent synthetic μ-opioid receptor (μOR) agonist, in conditions of low excitability *in vitro* when burstlets appeared at times bursts would have been expected or when preBötC rhythmic activity consisted solely of burstlets. We measured the frequency of inspiratory-related preBötC population activity, i.e., the intervals between successive bursts or burstlets, to assay opioidergic effects on rhythmogenesis and calculated the fraction of rhythmic preBötC events that were burstlets to assay changes in the pattern-generating threshold mechanism that converts burstlets into bursts. We conclude that DAMGO depresses inspiratory frequency by acting on burstlet producing preBötC Dbx1^+^ neurons. These data are consistent with our hypothesis of distinct rhythm and pattern generating mechanisms within the preBötC, with rhythmogenesis being mediated by burstlets and a separable burst-generating mechanism governing patterned preBötC output. Our results also inform potential approaches for combating opioid-induced depression of breathing.

## Results

### DAMGO slows preBötC rhythmic activity without affecting burstlet fraction

Systemic administration of opioids slows breathing *in vivo* and *in vitro* (Boom et al., 2012; Gray et al., 1999). We investigated how preBötC burstlets and bursts mediate these opioidergic effects in neonatal mouse tissue slices that contain the preBötC and the hypoglossal motor nucleus and nerve (XII) and generate a physiologically relevant motor output, i.e., inspiratory-related rhythmic activity. Under baseline conditions in artificial cerebrospinal fluid (ACSF) containing 9 mM K^+^ and 1.5 mM Ca^2+^ (“9/1.5”), preBötC population activity and XII activity consisted primarily of coincident, large amplitude rhythmic bursts (Fig. 1A; Smith et al., 1991). Subsaturating concentrations of DAMGO caused a dose-dependent decrease in the frequency (*f*) of rhythmic preBötC and XII bursts (Fig. 1A-C). 10 nM DAMGO significantly decreased preBötC *f* from 0.22 ± 0.05 Hz to 0.13 ± 0.04 Hz (p=0.01, n=6; Fig. 1C). 30 nM DAMGO application further decreased preBötC *f* to 0.07 ± 0.04 Hz (p=0.0001, n=6; Fig. 1C), and 100 nM DAMGO led to complete cessation of XII activity and, in some cases, also blocked all preBötC activity (Fig. 1B).

**Figure 1.**
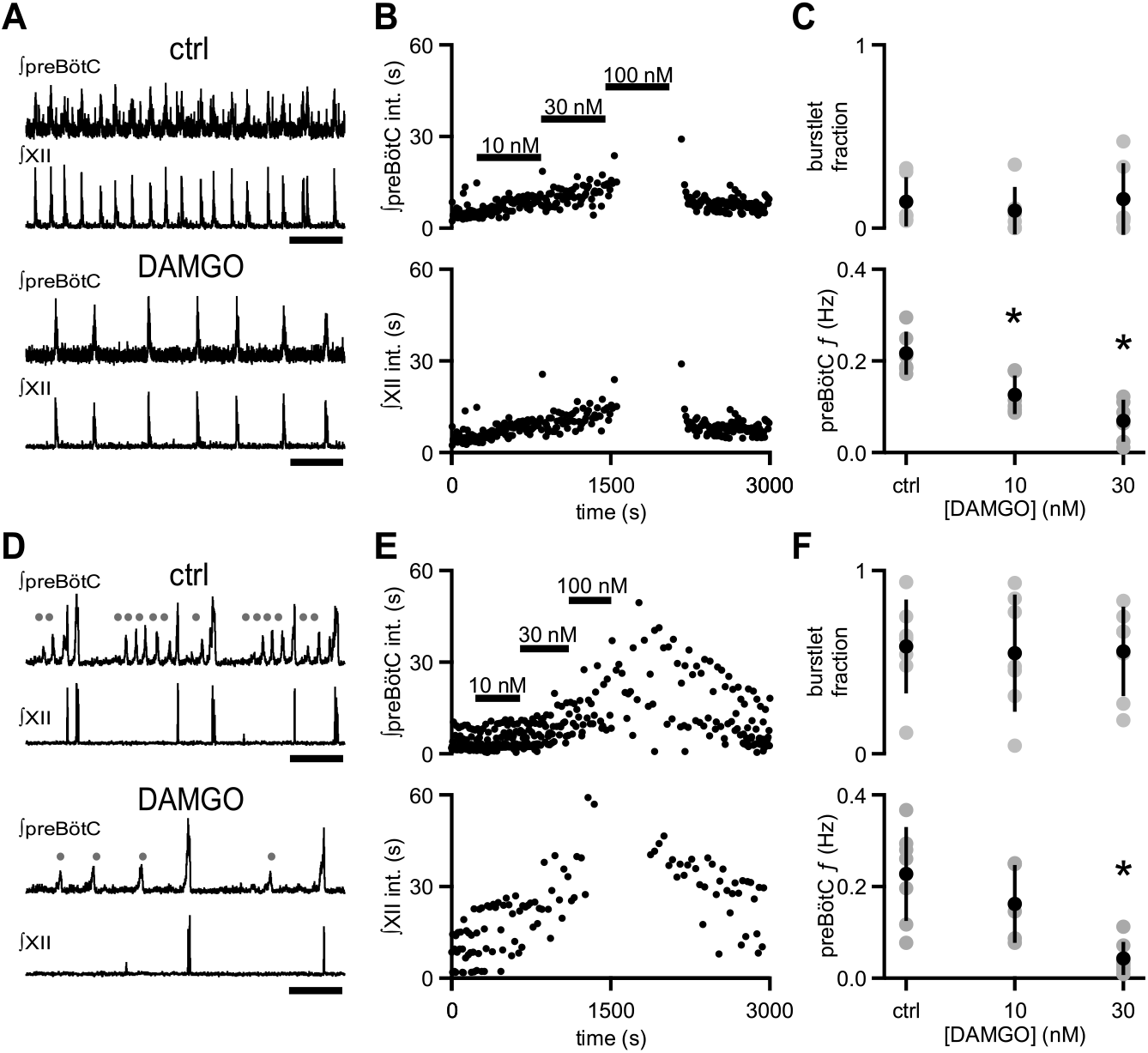
μOR activation depresses inspiratory frequency without affecting burstlet fraction. (A) Representative traces of ∫preBötC and ∫XII population activity in 9/1.5 ACSF control (ctrl) and 30 nM DAMGO. Scale bar, 10 s. (B) Representative time course of an experiment where increasing concentrations of DAMGO were bath-applied in 9/1.5 ACSF. Interevent intervals (int.) in preBötC (top) and XII (bottom) are plotted. In this experiment, 100 nM DAMGO led to cessation of rhythm. (C) Average burstlet fraction and preBötC *f* in 9/1.5 ACSF control condition (ctrl) and 10 nM and 30 nM DAMGO. Increasing concentrations of DAMGO did not significantly affect burstlet fraction whereas preBötC *f* was significantly lower in 10 nM and 30 nM DAMGO. *, p < 0.05, One-way ANOVA, post-hoc Tukey test, n=6. (D) Representative traces of ∫preBötC and ∫XII population activity in 3/1 ACSF control (ctrl) and 30 nM DAMGO. Both small amplitude burstlets (•), which did not produce XII activity, and large amplitude bursts, which generated XII bursts, were observed. Scale bar, 10 s. (E) Representative time course of an experiment, where increasing concentrations of DAMGO were bath-applied in 3/1 ACSF. Interevent intervals in preBötC (top) and XII (bottom) are plotted. In this experiment, 100 nM DAMGO led to cessation of XII activity while preBötC rhythm persisted. (F) Average burstlet fraction and *f* in 3/1 ACSF control condition (ctrl) and 10 nM and 30 nM DAMGO. Increasing concentrations of DAMGO did not significantly affect burstlet fraction whereas preBötC *f* was significantly lower in 30 nM DAMGO. *, p < 0.05, One-way ANOVA, post-hoc Tukey test, n=7.

We previously showed that changes in XII *f* induced by decreases in extracellular K^+^ could be caused by a reversion of preBötC bursts to preBötC burstlets (Kam et al., 2013a). This was not the case for the effects of DAMGO on *f*, as the fraction of total preBötC events that were burstlets (burstlet fraction), which was very low in control conditions (0.14 ± 0.14), was not significantly different from that in the presence of 30 or 100 nM DAMGO (p=0.8, n=6; Fig. 1A, C). Moreover, the amplitude of preBötC bursts (0.60 ± 0.10 a.u.) was not significantly different between control conditions without DAMGO and when 10 nM (0.61 ± 0.13 a.u.) or 30 nM DAMGO (0.56 ± 0.19 a.u.) was added to the bath (p=0.8, n=6).

To determine whether DAMGO could affect the conversion of burstlets into bursts when the preBötC was operating in a different dynamic mode, we decreased ACSF K^+^ and Ca^2+^ concentrations (Kam et al., 2013a). In 3 mM K^+^ and 1 mM Ca^2+^ (“3/1”), preBötC activity was a mixed pattern of burstlets and bursts (Fig. 1D, upper traces). While the burstlet fraction under these conditions (0.58 ± 0.26) was significantly higher than that in 9/1.5 ACSF (0.14 ± 0.14; p=0.003, n= 6 for 9/1.5, n=7 for 3/1), preBötC *f* was similar (0.23 ± 0.10 Hz in 3/1 vs. 0.22 ± 0.05 Hz in 9/1.5; p=0.8, n= 6 for 9/1.5, n=7 for 3/1; Fig. 1C, F). XII *f* in 3/1 ACSF (0.08 ± 0.05 Hz), on the other hand, was significantly lower than XII *f* in 9/1.5 ACSF (0.2 ± 0.06 Hz; p=0.002, n= 6 for 9/1.5, n=7 for 3/1; Fig. 1B, E), a consequence of the increased fraction of preBötC burstlets that were not transmitted to XII output.

In 3/1 ACSF, DAMGO decreased preBötC *f* in a dose-dependent manner, similar to its effects in 9/1.5 ACSF. Addition of 30 nM DAMGO significantly reduced preBötC *f* from 0.22 ± 0.1 Hz to 0.04 ± 0.04 Hz (p=0.001, n=7; Fig. 1D-F). This reduction was not significantly different from the effects of 30 nM DAMGO in 9/1.5 ACSF (p=0.2, n=6 for 9/1.5, n=7 for 3/1; Fig. 1C, F). Higher concentrations of DAMGO (≥100 nM) frequently blocked all preBötC population activity. Burstlet and burst amplitudes in 3/1 ACSF (burstlet: 0.38 ± 0.05 a.u.; burst: 0.73 ± 0.08 a.u.) did not change significantly when 10 nM DAMGO (burstlet: 0.36 ± 0.04 a.u.; burst: 0.78 ± 0.10 a.u.) or 30 nM DAMGO (burstlet: 0.34 ± 0.11 a.u.; burst: 0.65 ± 0.21 a.u.) was bath-applied (burstlet: p=0.7, burst: p=0.3, n=7), and the burstlet fraction was again unaffected by DAMGO (p=1.0, n=7; Fig. 1F).

To determine whether opioids specifically affected rhythmogenic activity independent of burst production, we isolated burstlets using low concentrations of Cd^2+^ (6-25 μM; Kam et al., 2013a), which significantly increased the burstlet fraction from 0.42 ± 0.30 to 0.78 ± 0.20 (p=0.02, n=8) without significantly changing preBötC *f* or amplitude (*f*: p=0.7, amplitude: p=0.06, n=8; Fig. 2A, B). The decreases in excitability and synaptic efficacy caused by Cd^2+^ (Kam et al., 2013a; Lu et al., 2007) resulted in an increased sensitivity to DAMGO. Bath application of 1 nM DAMGO on this burstlet only rhythm was sufficient to abolish preBötC rhythmic activity completely in 7 of 8 slices. Concentrations of DAMGO below that causing rhythmic cessation (0.01-10 nM) slowed preBötC *f* significantly from 0.28 ± 0.09 Hz to 0.18 ± 0.07 Hz (p=0.001, n=8; Fig. 2A, B). This relative decrease in preBötC *f* (0.66 ± 0.16) was not significantly different from the effect of 10 nM DAMGO on preBötC *f* in 3/1 ACSF with no Cd^2+^ (0.63 ± 0.16; p=0.8, n=5 for 3/1, n=8 for Cd^2+^). However, burstlet amplitudes in the presence of both Cd^2+^ and DAMGO (0.29 ± 0.08) were significantly reduced compared to Cd^2+^ alone (0.35 ± 0.09; p=0.003, n=8; Fig. 2A, B).

**Figure 2.**
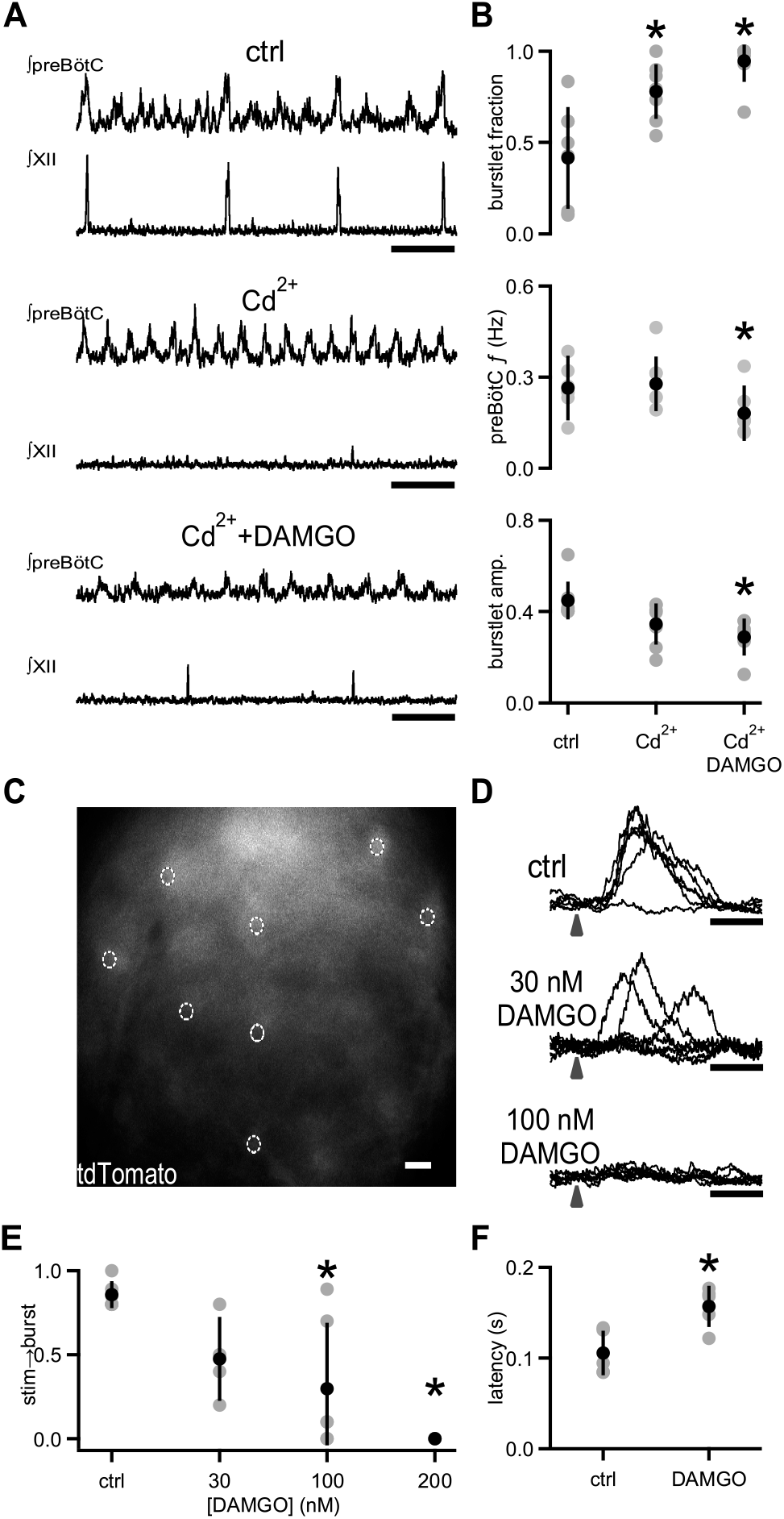
DAMGO specifically modulates preBötC rhythmogenesis. (A) Representative traces of ∫preBötC and ∫XII population activity in 3/1 ACSF control (ctrl), 8 μM Cd^2+^, and 8 μM Cd^2+^ + 10 nM DAMGO. In 3/1 ACSF (top), rhythmic activity consisted of both burstlets and bursts. After Cd^2+^ application (middle), preBötC bursts were no longer observed, and XII activity was abolished, leaving only a preBötC burstlet rhythm with a *f* similar to control. With addition of DAMGO (bottom), burstlet *f* and amplitude decreased. Scale bar, 5 s. (B) Average burstlet fraction, preBötC *f*, and burstlet amplitude (amp.) in 3/1 ACSF control (ctrl), 6-25 μM Cd^2+^, and 6-25 μM Cd^2+^ + 0.1-10 nM DAMGO. Addition of Cd^2+^ significantly increased the burstlet fraction while preBötC *f* and burstlet amplitudes were unchanged. Bath application of DAMGO significantly decreased preBötC *f* (consisting almost solely of burstlets) and burstlet amplitude. *, p < 0.05, One-way ANOVA, post-hoc Tukey test, n=8. (C) Representative image of tdTomato fluorescence in the preBötC of a medullary slice from a *Dbx1*^*cre*^;*Rosa26*^*tdTomato*^ mouse. Dashed circles represent holographic spots over targeted Dbx1^+^ neurons. Scale bar, 20 μm. (D) Overlays of representative traces of ∫preBötC from successive laser stimulations of the 8 Dbx1^+^ neurons shown in (C) in control 9/1.5 ACSF (ctrl), 30 nM DAMGO, and 100 nM DAMGO. Arrow represents time of laser stimulation. In control conditions, a failure was seen, but burst initiation was otherwise reliably successful. Bursts were consistently triggered after a ~100 ms latency. In 30 nM DAMGO, failures were more frequent when the same stimulation parameters were used, and the latency between laser stimulation and burst initiation was longer and more variable. In 100 nM DAMGO, bursts were no longer triggered with the same stimulation parameters. Scale bar, 200 ms. (E) Average success rate (stim→burst) for burst initiation in control 9/1.5 ACSF (ctrl) conditions and in increasing concentrations of DAMGO using entraining stimuli that elicits >80% success in 9/1.5 ACSF. To entrain the rhythm, 5-10 Dbx1^+^ neurons were stimulated 8 - 20 times every 3-6 s. DAMGO produced a dose-dependent decrease in the success rate for triggering bursts with a significant reduction in success between control and at 100 nM and 200 nM DAMGO. *, p < 0.05, One-way ANOVA, post-hoc Tukey test, n=6. (F) Average latency between laser stimulation and burst initiation when bursts were successfully triggered during entraining stimuli was increased in 30 nM DAMGO compared to control 9/1.5 ACSF (ctrl). *, p<0.05, Student’s t-test, n=5.

Depression of preBötC burstlet *f* by μOR activation points to DAMGO modulating preBötC rhythmogenic mechanisms. We previously hypothesized that the emergent rhythmogenic processes that produce burstlets manifested as the latency in ectopic burst initiation following targeted excitation of 4-9 preBötC inspiratory neurons, whose molecular identity was unspecified (Kam et al., 2013b). Here, in slices from Dbx1^+^ reporter (*Dbx1*^*cre*^;*Rosa26*^*tdTomato*^) mice, we targeted preBötC Dbx1^+^ neurons for holographic photostimulation to determine if opioids also modulate this rhythmogenesis-related process (Fig. 2C-F). In control 9/1.5 ACSF, ectopic bursts were reliably elicited when 5-10 Dbx1^+^ neurons were targeted (probability of success: 0.86 ± 0.08, n=6; Fig. 2D). When a set of neurons that reliably resulted in bursts in response to photostimulation in control conditions were then excited in 30 nM DAMGO, the probability of eliciting a burst was reduced to 0.48 ± 0.25 (n=4; Fig. 2D, E). In 100 nM DAMGO, the success rate decreased significantly as stimulation of the threshold set that had previously reliably triggered bursts was further compromised and, in some cases, failed to trigger a single burst (p=0.006, n=6; Fig. 2D, E). In 200 nM DAMGO, stimulation was unsuccessful at eliciting bursts (p=0.0003, n=6; Fig. 2E). Importantly, the latency between laser stimulation and burst initiation in the cases when bursts were triggered in DAMGO (157 ± 22 ms) was significantly increased compared to that in baseline conditions (105 ± 24 ms; p=0.002, n=5; Fig. 2F).

### μOR-expressing Dbx1^+^ neurons mediate the rhythmogenic effects of opioids

Which preBötC neurons does DAMGO target to depress inspiratory rhythm? We hypothesized that DAMGO was acting on preBötC Dbx1^+^ neurons, which are essential for respiratory rhythm generation (Bouvier et al., 2010; Gray et al., 2010; Wang et al., 2014) and contain μOR mRNA (Hayes et al., 2017). We first sought to determine whether these neurons express μOR protein using immunohistochemistry. In *Dbx1*^*Cre*^;*Rosa26*^*tdTomato*^ mice, μOR protein expression overlapped with the spatial distribution of tdTomato-expressing Dbx1^+^ neurons in preBötC and the adjacent intermediate band of the reticular formation (IRt; Fig. 3A) and was localized to individual preBötC Dbx1^+^ neurons (Fig. 3B-C).

**Figure 3.**
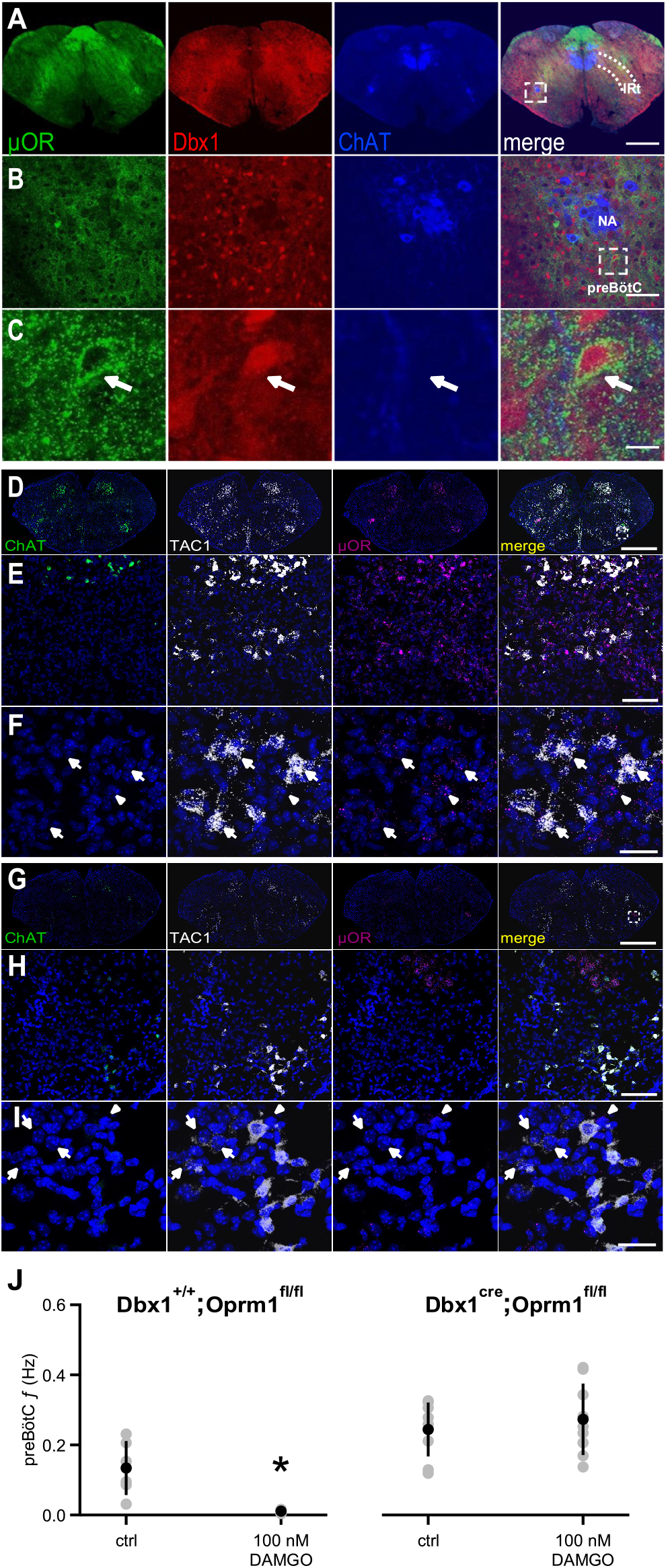
μORs expressed in Dbx1^+^ neurons mediate DAMGO inhibition of preBötC rhythmic activities. (A) Confocal images of brainstem sections of a *Dbx1*^*Cre*^;*Rosa26*^*tdTomato*^ mouse pup (P2) double-stained with antibodies to ChAT (blue) and μOR (green). Coronal sections showing μOR expression in the preBötC and the Intermediate band of the Reticular Formation (IRt) and colocalization with Dbx1^+^ neurons in the preBötC. Scale bar, 500 μm. (B) Higher magnification of nucleus ambiguus (NA) and preBötC from the boxed region in A. Scale bar, 50 μm. (C) Expanded images of the boxed preBötC region highlighted in B. The arrow shows a preBötC Dbx1^+^ neuron with abundant punctate μOR expression on the cell body. Scale bar, 10 μm. (D) Confocal image of a medullary brainstem section at the level of the preBötC from a control P2 *Dbx1*^*+/+*^; *Oprm1*^*fl/fl*^ mouse processed with RNAScope. Probes for μOR (magenta), TAC1 (white), and acetylcholine esterase (ChAT; green) were used, and the tissue was counterstained with DAPI (blue). A box marks the location of preBötC. Scale bar, 1 mm. (E) Higher magnification of the boxed preBötC region in D. μOR and TAC1 were highly expressed in this region. Scale bar, 100 μm. (F) Higher magnification images of μOR-expressing neurons with (arrows) and without (arrowhead) TAC1 expression from E. Colocalization of TAC1 and multiple μOR puncta was observed in several neurons as well as abundant μOR puncta in neurons that did not express TAC1. Scale bar, 40 μm. (G) Confocal image of a medullary brainstem section at the level of the preBötC from a P2 *Dbx1*^*cre/cre*^; *Oprm1*^*fl/fl*^ mouse processed with RNAScope. Probes for μOR (magenta), TAC1 (white), and acetylcholine esterase (ChAT; green) were used, and the tissue was counterstained with DAPI (blue). In the preBötC of these mice, the density of μOR puncta was low. A box marks the location of preBötC. Scale bar, 1 mm. (H) Higher magnification of the boxed preBötC region in G. The density of μOR puncta was low, and μOR expression was not associated with TAC1 expression. Scale bar, 100 μm. (I) Higher magnification images of TAC1-expressing neurons from H with few to no μOR puncta (arrows) as well as μOR-expressing neurons without TAC1 expression (arrowhead). Scale bar, 40 μm. (J) Average *f* of preBötC activity in control 9/1.5 conditions (ctrl) and 100 nM DAMGO in *Dbx1*^*Cre*^; *Oprm1*^*fl/fl*^ mice (right), where μORs are selectively deleted in Dbx1^+^ neurons, and their *Dbx1*^*+/+*^;*Oprm1*^*fl/fl*^ littermate controls with preserved μOR expression (left). Whereas 100 nM DAMGO reduced preBötC *f* significantly in *Dbx1*^*+/+*^;*Oprm1*^*fl/fl*^ control mice, the same concentration had no effect on *f* when μORs were deleted. *, p<0.05, Student’s t-test, n=6 for *Dbx1*^*+/+*^;*Oprm1*^*fl/fl*^, n=9 for *Dbx1*^*Cre*^;*Oprm1*^*fl/fl*^.

To test whether μORs in Dbx1^+^ neurons mediated DAMGO inhibition, we used a *Dbx1*^*Cre*^;*Oprm1*^*fl/fl*^ mouse line (Weibel et al., 2013), in which μORs are genetically deleted in Dbx1^+^ neurons. We first examined whether preBötC μOR expression was knocked down in P2 *Dbx1*^*Cre*^;*Oprm1*^*fl/fl*^ mice compared to their littermate controls (Fig. 3D-I). *Dbx1* is not expressed postnatally (Bouvier et al., 2010; Gray et al., 2010), and NK1R protein is predominantly found in neuronal processes (Tan et al., 2010) while NK1R mRNA is sparsely expressed in somata (Fig. S1). We therefore localized preBötC using tachykinin precursor peptide (TAC1) expression, which colocalizes with NK1R, ventral to nucleus ambiguus (NA) as a marker for preBötC (Fig. S1; Hayes et al., 2017). To measure μOR and TAC1 expression, we performed fluorescence *in situ* hybridization, which avoids the non-specific labeling that may occur with some μOR antibodies (Schmidt et al., 2013) and allows localization of TAC1 expression to somata (Fig. S1). Abundant (>15 puncta) TAC1 mRNA was found in larger neurons confined to the preBötC (somatic diameter: 27.08 ± 4.3 μm, n=121). Choline acetyl transferase (ChAT) mRNA probes were used to mark NA. In *Dbx1*^*+/+*^; *Oprm1*^*fl/fl*^ control mice, μOR mRNA puncta colocalized with TAC1 and NK1R mRNA in the preBötC (Figs. 3D-F, S1), recapitulating the immunohistochemical distribution of Dbx1^+^ neurons in the Dbx1^+^ reporter mice (Fig. 3A-C). Compared with μOR expression in *Dbx1*^*+/+*^; *Oprm1*^*fl/fl*^ mice (Fig. 3D-F), μOR expression in *Dbx1*^*Cre*^; *Oprm1*^*fl/fl*^ mice was reduced in regions where Dbx1^+^ neurons are normally found (Fig. 3D, G) and significantly reduced in TAC1 positive preBötC neurons (p=0.01, *Dbx1*^*+/+*^; *Oprm1*^*fl/fl*^: H-score = 139.07 ± 39.33, n = 5; *Dbx1*^*Cre*^; *Oprm1*^*fl/fl*^: H-score= 47.98 ± 19.77, n=3; Fig. 3G-I).

Having confirmed a decrease in μOR expression in preBötC in *Dbx1*^*Cre*^; *Oprm1*^*fl/fl*^ mice, we determined whether the functional effects of DAMGO on preBötC rhythm were similarly affected. In rhythmic tissue slices from *Dbx1*^*Cre*^; *Oprm1*^*fl/fl*^ mice bathed in 9/1.5 ACSF, 100 nM DAMGO had little effect on preBötC *f* (p=0.2, n=9; Figs. 3J, S2). In contrast, bath-applied DAMGO in *Dbx1*^*+/+*^; *Oprm1*^*fl/fl*^ cre-negative littermates reduced preBötC *f* significantly (p=0.01, n=6; Fig. 3J).

### DAMGO-mediated depression is diminished by Substance P, but not blockade of inhibitory synaptic transmission

The coexpression of μOR and NK1R in preBötC Dbx1^+^ neurons raises the possibility that NK1R activation, which increases respiratory frequency in wild type rodents (Gray et al., 1999; Yeh et al., 2017), could be effective in reducing the depressant effects of DAMGO. When applied together in 3/1 ACSF, 30 nM DAMGO was much less effective in reducing rhythmic activity in the presence of the NK1R agonist Substance P (SP; 500 nM), and preBötC *f* was not significantly different from that in control 3/1 ACSF conditions, i.e., in the absence of SP and DAMGO (p=0.09, n=6; Fig. 4A,C). We next tested whether altering preBötC excitability by blockade of inhibitory synaptic transmission using a cocktail of the GABA_A_ receptor antagonist picrotoxin and the glycine receptor antagonist strychnine (picrotoxin/strychnine; Del Negro et al., 2009; Ren and Greer, 2006; Shao and Feldman, 1997) would similarly block DAMGO-mediated depression. In contrast to SP, picrotoxin/strychnine (100 μM/1 μM) did not reduce the effects on preBötC *f* of 30 nM DAMGO compared to control conditions in either 9/1.5 or 3/1 ACSF (p=0.2, n=6 for 9/1.5, n=7 for 3/1, n=13 for picrotoxin/strychnine; Fig. 4B, C). Indeed, preBötC *f* when SP was applied with 30 nM DAMGO was greater than that in 9/1.5 ACSF, with or without picrotoxin/strychnine, or 3/1 ACSF in the presence of DAMGO (p=0.001, n=6 for 9/1.5, n=7 for 3/1, n=12 for picrotoxin/strychnine, n=6 for SP; Fig. 4C).

**Figure 4.**
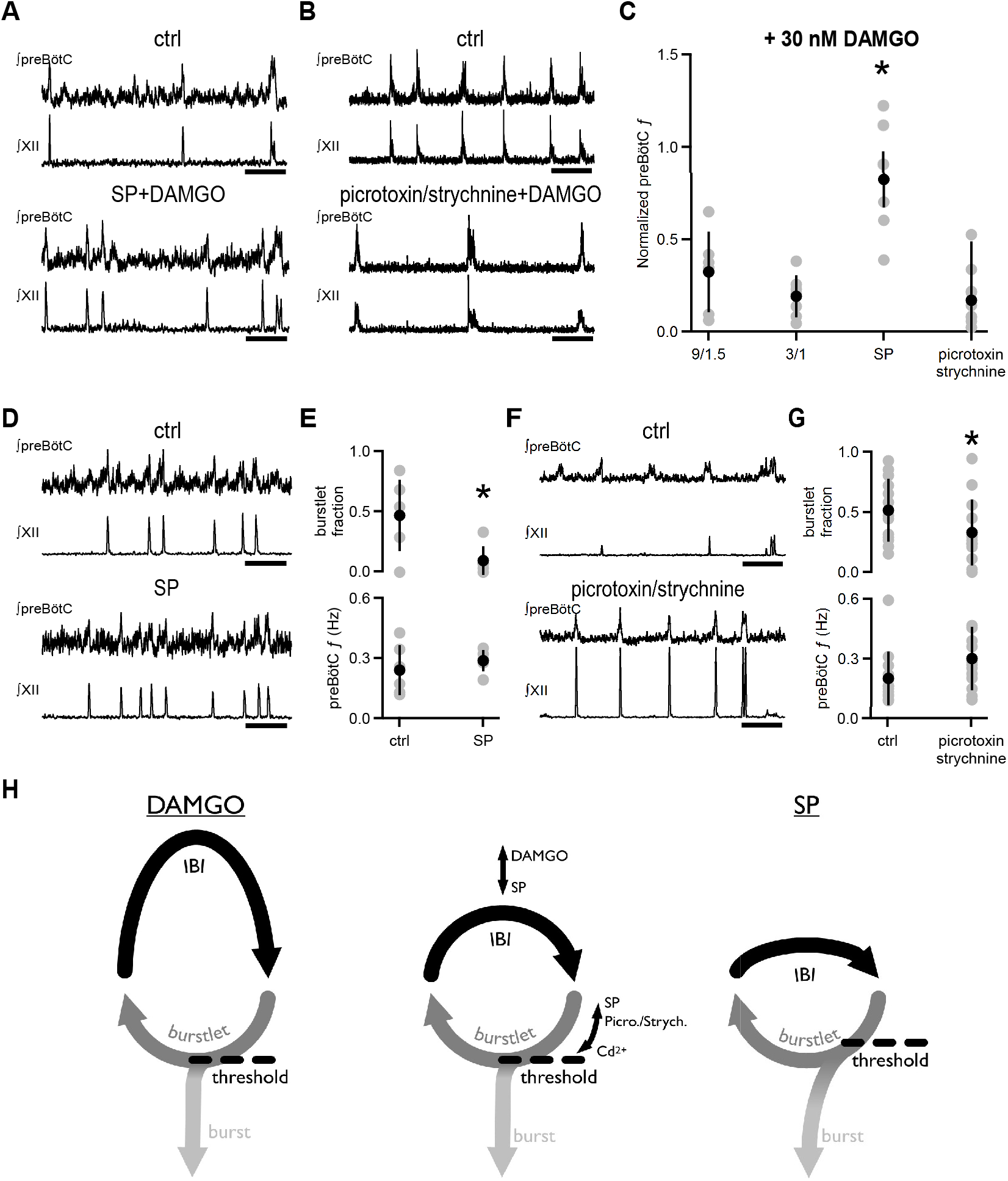
Substance P, but not blockade of inhibitory synaptic transmission, relieves DAMGO-mediated depression. (A) Representative traces of ∫preBötC and ∫XII population activity in 3/1 ACSF control (ctrl) and 500 nM SP and 30 nM DAMGO. SP mitigated the effects of DAMGO, and preBötC activity resembled that in control conditions. Scale bar, 10 s. (B) Representative traces of ∫preBötC and ∫XII population activity in 3/1 ACSF control (ctrl) and 100 μM picrotoxin/1 μM strychnine and 30 nM DAMGO. The addition of picrotoxin/strychnine following DAMGO application did not recover preBötC *f* to control levels. Scale bar, 10 s. (C) Average normalized *f* of preBötC activity when 30 nM DAMGO was added alone (9/1.5, 3/1) or following application of either 500 nM SP or 100 μM picrotoxin/1 μM strychnine with 30 nM DAMGO. For each experiment, values were normalized to preBötC *f* during baseline 9/1.5 or 3/1 ACSF in that experiment. When picrotoxin/strychnine were added with DAMGO, the relative change in *f* did not differ from the normalized *f* when DAMGO was applied alone in either 9/1.5 ACSF or 3/1 ACSF. The normalized *f* when SP was added with DAMGO was significantly higher than all other conditions. *, p < 0.05, One-way ANOVA, post-hoc Tukey test, n=6 for 9/1.5, n=7 for 3/1, n=12 for picrotoxin/strychnine, n=6 for SP. (D) Representative traces of ∫preBötC and ∫XII population activity in 3/1 ACSF control (ctrl) and 500 nM SP. In 3/1 ACSF, rhythmic activity consisted of both burstlets and bursts. After SP, *f* was unchanged, but more preBötC bursts were observed. Scale bar, 10 s. (E) Average burstlet fraction and *f* in 3/1 ACSF control condition (ctrl) and 500 nM SP. SP significantly decreased the burstlet fraction, but did not change preBötC *f*. *, p < 0.05, Student’s t-test, n=13. (F) Representative traces of ∫preBötC and ∫XII population activity in 3/1 ACSF control (ctrl) and 100 μM picrotoxin/1 μM strychnine. In 3/1 ACSF, rhythmic activity consisted of both burstlets and bursts. After picrotoxin/strychnine, *f* was unchanged; however, more preBötC bursts were observed, and XII bursts were larger. Scale bar, 10 s. (G) Average burstlet fraction and *f* in 3/1 ACSF control condition (ctrl) and 100 μM picrotoxin/1 μM strychnine. Picrotoxin/strychnine significantly decreased the burstlet fraction, but did not change preBötC *f*. *, p < 0.05, Student’s t-test, n=13. (H) Model for preBötC rhythm and pattern generation. In the absence of neuromodulation (center), rhythm generation is the alternation between burstlet activity and the silent interburstlet interval (IBI). Pattern generation in the preBötC occurs when burstlet activity reaches a threshold (dashed line) and triggers a burst that is then transmitted to downstream premotoneuronal and motoneuronal populations. These processes can be regulatedindependently by a variety of factors (double-headed arrows). Activation of μORs expressed in Dbx1^+^ neurons (left) slows the rhythm by decreasing excitability in Dbx1^+^ neurons, reducing synchrony, prolonging IBI, and reducing burstlet amplitude. SP (right) may mitigate the effects of μOR activation by increasing *f* or reducing the threshold for burst generation.

Are the differential efficacies of SP and blockade of inhibitory transmission expressed in their effects on baseline rhythmic activity? To determine how SP and blockade of inhibitory transmission affect rhythm and pattern generation, we tested the effects of SP and picrotoxin/strychnine on preBötC activity in 3/1 ACSF without DAMGO. Under these conditions, bath-applied SP (500 nM) significantly reduced the burstlet fraction from 0.47 ± 0.3 to 0.10 ± 0.12 (p=0.01, n=6), but, surprisingly, did not affect preBötC *f* (p=0.4, n=6; Fig. 4D, E). In slices from *Dbx1*^*Cre*^; *Oprm1*^*fl/fl*^ mice bathed in 9/1.5 ACSF, we confirmed that 500 nM SP significantly increased preBötC *f* (Fig. S2) and modulated preBötC *f* independent of μORs, which were deleted in Dbx1^+^ neurons in these slices. In 3/1 ACSF, bath-applied picrotoxin/strychnine (100 μM/1 μM) also significantly decreased burstlet fraction (p=0.0001, n=13) and did not alter preBötC *f* significantly (p=0.9, n=13; Fig. 4F, G).

## Discussion

Inspiratory rhythm generation must be both robust, to operate almost continuously throughout the lifetime of the animal, and labile, to adapt to changing metabolic and environmental conditions and coordinate with other behaviors. How the preBötC generates rhythm to meet these challenges has thus far defied a thorough understanding. We put forth a hypothesis that respiratory rhythmogenesis is driven not by strong bursts, but by preBötC burstlets (Fig. 4H; Kam et al., 2013a). Burstlets occur when spontaneous firing among preBötC neurons reaches a threshold for synchrony, and the time required for preBötC neurons to assemble and achieve this synchrony (percolation) determines the period of the rhythm (Kam et al., 2013a; S.A. and J.L.F., in preparation).

A strong prediction of this hypothesis is that manipulations that depress burst frequency should also slow burstlet frequency. Indeed, we found that DAMGO specifically slowed preBötC *f* without affecting burstlet fraction when the rhythm: i) was composed of mostly bursts; ii) was a mixed pattern of bursts and burstlets; or iii) was mostly (or all) burstlets (Fig. 4H). Additionally, DAMGO prolonged the latency between photostimulation and burst generation when threshold sets of ≤10 neurons were excited. The congruent effects of DAMGO on both burstlet *f* and photostimulation latency: i) is consistent with our hypothesis that both are products of percolation of activity within the preBötC microcircuit, a critical process for rhythmogenesis (Kam et al., 2013a; Kam et al., 2013b), and ii) suggests that DAMGO directly affects rhythmogenic mechanisms.

Consistent with our functional studies, we found abundant μOR expression in preBötC Dbx1^+^ neurons that form the core circuitry for inspiratory rhythmogenesis (Wang et al., 2014). We also showed colocalization of μOR with NK1R in some preBötC neurons as well as with a partially overlapping preBötC subpopulation expressing TAC1 that may be one endogenous source of SP (Liu et al., 2004). When μORs were genetically deleted in Dbx1^+^ neurons in *Dbx1*^*Cre*^; *Oprm1*^*fl/fl*^ mice, the potent rhythmic slowing effect of DAMGO in slices was no longer present. These data suggest that μORs in preBötC Dbx1^+^ neurons mediate DAMGO inhibition of rhythmicity.

How could μOR activation affect the percolation of activity in the preBötC that we hypothesize underlies rhythmogenesis and transform, i.e., slow down, the dynamics of the Dbx1^+^ preBötC network? μORs can be localized postsynaptically on dendrites and cell bodies, where they regulate neuronal excitability and transduce receptor activation to downstream signal transduction pathways; μORs are also found presynaptically on axon terminals where they inhibit neurotransmitter release via activation of K^+^ conductances and/or inhibition of Ca^2+^ conductances (Le Merrer et al., 2009; Mansour et al., 1988; Montandon et al., 2016; Williams et al., 2001). We suggest that DAMGO-mediated hyperpolarization and/or decreases in synaptic probability of release in and among μOR-expressing preBötC Dbx1^+^ neurons reduce spontaneous firing rate, weaken the propagation of action potentials to postsynaptic neurons, and increase the number of inputs required to reach action potential threshold. These effects reduce the moment-to-moment probability that a critical number of neurons will be simultaneously active and prolong the time required for preBötC neurons to increase their firing, achieve synchrony, and generate burstlets (S.A. and J.L.F., in preparation).

The pattern-generating mechanism that converts burstlets to bursts may, by analogy, require a higher threshold of activity and synchrony be reached via independently regulated, distinct mechanisms, such as the activation of persistent inward conductances (Picardo et al., 2019). Through this threshold mechanism, SP and blockade of inhibitory synaptic transmission increase, whereas Cd^2+^ decreases, the fraction of preBötC events that are burstlets, without altering frequency of these events (Fig. 4H). Interestingly, the depressant effects of DAMGO were reduced by SP, but not inhibitory blockade. Inhibitory synaptic activity in preBötC is dispensable for generation of breathing rhythm *in vivo* and *in vitro* (Janczewski et al., 2013; Shao and Feldman, 1997) and has variable effects on excitability (Ren and Greer, 2006), which may account for the inability of picrotoxin and strychnine to mitigate DAMGO-induced depression. Alternatively, inhibition may act elsewhere in the microcircuit or upstream of opioidergic signaling pathways within Dbx1^+^ neurons.

While inhibitory blockade specifically affected pattern generation and was unable to rescue DAMGO-mediated depression, NK1R activation seems capable of modulating both rhythm and pattern in a state-dependent manner that can offset the effects of DAMGO (Fig. 4H). A ceiling effect, i.e., preBötC *f* being near or at maximal, may have limited the rhythmogenic effect of SP in 3/1 ACSF alone, but did reveal a pattern generating effect on burstlet fraction. Consistent with SP effects in 9/1.5 ACSF (Gray et al., 1999; Yeh et al., 2017), co-application of opioids and SP demonstrated a role for NK1R activation in modulating rhythm that is likely mediated by intracellular interactions in preBötC Dbx1^+^ neurons that coexpress μOR and NK1R. The μOR expression and DAMGO sensitivity of NK1R-expressing neurons are well-documented (Gray et al., 1999; Montandon et al., 2011), and G_s_-protein coupled signaling of NK1R opposes the G_i/o_ coupled signaling of μORs (Gray et al., 1999; Johnson et al., 1996). However, the interaction of NK1R and μOR can be complicated. Opioids may inhibit SP release, and SP potentiates the antinociceptive effects of morphine through release of endogenous opioids (Fukazawa et al., 2007; Kream et al., 1993). SP may also act directly on μORs as a weak agonist since naloxone, the μOR antagonist, blocks SP excitation in some neurons (Davies and Dray, 1977). Nonetheless, we predict that only neuropeptides and neuromodulators, whose receptors are coexpressed with μORs in rhythmogenic preBötC neurons, like NK1Rs, are able to modulate μOR signaling intracellularly to mitigate DAMGO-mediated depression.

The interaction between excitatory neuromodulatory systems with opiates in respiratory neural circuits is relevant for addressing the depressive effects of opiates on breathing. Opioids are the most commonly prescribed drugs for severe acute and chronic pain and play an important role in palliative care. However, with a high potential for addiction, opiate overprescription and abuse has created a pressing public health crisis that affects millions (CBHSQ, 2018). Morbidity and mortality from opioid addiction and overdose is largely a result of opioid-induced depression of breathing (Jaffe & Martin, 1990). Opiates given systemically will act on opioid receptors throughout the nervous system (Kibaly et al., 2019), and μORs are expressed in various brain structures regulating breathing (Gray et al., 1999; Manzke et al., 2003; Phillips et al., 2012; Pokorski and Lahiri, 1981; Prkic et al., 2012; Zhang et al., 2007; Zhang et al., 2011). While depression of breathing is unlikely to be dependent on actions at a single site (Lalley et al., 2014; Montandon and Horner, 2014; Stucke et al., 2015), our data is consistent with compelling *in vitro* and *in vivo* evidence that the preBötC is the most sensitive site to μOR agonists and that it mediates respiratory frequency depression by opioids (Ballanyi and Ruangkittisakul, 2009; Janczewski et al., 2002; Montandon and Horner, 2014; Montandon et al., 2011).

In summary, the depressive effect of the μOR agonist DAMGO on preBötC burstlet frequency, in the absence of a change in burstlet fraction, supports a primary role for burstlets in respiratory rhythm generation and is consistent with the hypothesis that rhythm and pattern generating mechanisms within the preBötC are separable. We propose that having distinct mechanisms generating burstlets and bursts provides functional substrates for targeted modulation of the rhythmic timing and/or pattern of breathing by other brain areas and neuromodulators. This organization enables the dynamic temporal and pattern lability characteristic of breathing essential for adaptation to changing metabolic demands and for coordination with other respiratory-related and orofacial behaviors. A better understanding of the mechanisms of μOR actions in respiratory neural circuits has important implications for developing stimulants that excite breathing via non-opioidergic pathways (van der Schier et al., 2014) and drugs that reverse opioid actions on preBötC Dbx1^+^ neurons to maintain breathing without affecting analgesia.

## Acknowledgements

The authors thank Grace Li for excellent technical work. *Oprm1*^*fl/fl*^ mice were generously provided by Dr. Wendy Weibel, Dr. Chris Evans, and the Animal Breeding Core (NIDA P50 Center Grant, DA005010). This work was supported by National Institutes of Health grants NS07221, HL135779, and NS097492; and an international postdoctoral grant from the Swedish Research Council (Vetenskapsrådet).

## Author contributions

X.S., J.L.F., and K.K. designed the research; X.S., C.T.P., N.H., X.M.S., M.G., S.H., and K.K. performed the research; X.S., C.T.P. and K.K. analyzed the data; X.S., C.T.P., J.L.F., and K.K. wrote the paper.

## Experimental Procedures

Experimental procedures were carried out in accordance with the United States Public Health Service and Institute for Laboratory Animal Research Guide for the Care and Use of Laboratory Animals. All protocols were approved by University of California Animal Research Committee and the Rosalind Franklin University of Medicine and Science Institutional Animal Care and Use Committee. Every effort was made to minimize pain and discomfort, as well as the number of animals.

### Transgenic mice

*Dbx1*^*Cre*^ mice (Bielle et al., 2005) were crossed with either *Rosa26*^*tdTomato*^ (JAX Stock No. 007908) or *Oprm1*^*fl/fl*^ (Weibel et al., 2013) mouse lines. *Dbx1*^*Cre*^;*Rosa26*^*tdTomato*^ were used to visualize Dbx1^+^ neurons, and *Dbx1*^*Cre*^;*Oprm1*^*fl/fl*^ mice were used to delete μORs selectively in Dbx1^+^ neurons using cre-lox recombination. Animal genotypes were verified via real time polymerase chain reaction using primers specific for the *Oprm1*^*fl/fl*^ conditional allele or cre recombinase (Pierani et al., 2001; Rose et al., 2009; Weibel et al., 2013) or by direct visualization of fluorescent reporter.

### Immunohistochemistry and confocal imaging

*Dbx1*^*Cre*^;*Rosa26*^*tdTomato*^ and *Dbx1*^*Cre*^;*Oprm1*^*fl/fl*^ mice were anesthetized at P0-4 by inhalation of isoflurane. The brainstem was isolated from the pons to the rostral cervical spinal cord, fixed in 4% paraformaldehyde, and cryoprotected in 30% sucrose before sectioned at 70mm thickness using CRYOSTAR NX70 (Thermo Scientific). Tissue sections were washed in PBS, and incubated in primary antibody at least overnight, followed by incubation in secondary antibody for 2 hrs at room temperature, and cover-slipped in Vectashield with or without DAPI. Primary antibodies used included rabbit anti-μOR (1:400), goat anti-ChAT (1:400, SCBT); Secondary antibodies were species specific (1:250) and conjugated to Alexa 488, Rhodamine or Cy5 (Invitrogen or Jackson ImmunoResearch). Sections were imaged using confocal laser scanning microscopy. Confocal images were captured on a Zeiss LSM 710 Meta confocal microscope implemented on an upright Axioplan 2 microscope. Zen software was used to capture images. Confocal images were analyzed with ImageJ, and the final figures were composed in Adobe Photoshop and/or Microsoft Powerpoint.

### In situ hybridization

As described above, mice were anesthetized at P0-4 by inhalation of isoflurane. The isolated brainstem was rapidly frozen in dry ice and stored at −80°C until sectioning. Transverse sections (14-20 μm) were cut on a cryostat (CryoStar NX70, Thermo Scientific) and mounted on SuperFrost Plus (Fisher Scientific) slides. Slides were processed for fluorescence *in situ* hybridization of three target RNAs simultaneously according to manufacturer protocols for RNAScope^®^ (Advanced Cell Diagnostics). Briefly, fresh frozen tissue samples were postfixed in 10% neutral buffered formalin, washed, and dehydrated in sequential concentrations of ethanol (50, 70 and 100%) at RT. Samples were treated with protease IV, then incubated for two hours at 40°C in the presence of target probes to allow for hybridization. The following target probes were used: Mm-Oprm1, Mm-Tac1, and Mm-ChAT. A series of three amplification steps was necessary to provide substrates for target fluorophore labeling (ChAT conjugated to fluorescein (1:1200), Oprm1 to Cy5, (1:1200) and Tac1 to Cy3 (1:1500) (Jackson ImmunoResearch). After labeling, samples were counterstained with DAPI. ProLong^®^ Gold (Invitrogen), was applied to each slide prior to cover slipping. Images were acquired on a confocal laser scanning microscope (LSM710 META, Zeiss). High-resolution z-stack confocal images were taken at 0.3 μm intervals. Cell-by-cell μOR expression profiles were quantified in selected regions of interest (TAC1 positive neurons in preBötC) according to a five-grade scoring system recommended by the manufacturer (ACD score; 0, no staining; 1, 1-3 dots/cell; 2, 4-9 dots/cell; 3, 10-15 dots/cell; 4, >15 dots/cell), where a dot (>0.60 μm) transcript. A Histo score (H-score) was then calculated from the data, using image-based software analysis (FIJI):

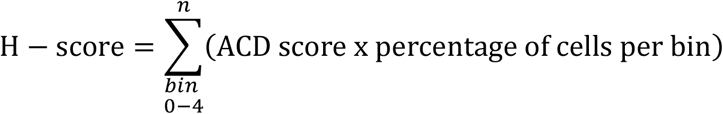

### Slice preparation and electrophysiology

Neonatal C57BL/6 or mutant mice (P0-5) of either sex were used for cutting brainstem medullary transverse slices (550-600 μm thick) containing the rhythm-generating preBötC, XII inspiratory motoneurons, and premotoneurons linking these populations (Koizumi et al., 2008; Smith et al., 1991). To obtain slices with the preBötC at the surface, the rostral cut was made above the first set of XII nerve rootlets at the level of the dorsomedial cell column and principal lateral loop of the inferior olive, and the caudal cut captured the obex (Ruangkittisakul et al., 2011). Slice cutting artificial cerebrospinal fluid (ACSF) solution was composed of (in mM): 124 NaCl, 3 KCl, 1.5 CaCl_2_, 1 MgSO_4_, 25 NaHCO_3_, 0.5 NaH_2_PO_4_, and 30 D-glucose, equilibrated with 95% O_2_ and 5% CO_2_, 27°C, pH 7.4. Slices were placed rostral side up in a chamber and perfused at 2-3 ml/min with 28-30°C recording ACSF. Baseline recording ACSF consisted of either the cutting ACSF solution with K^+^ raised to 9 mM and Ca^2+^ maintained at 1.5 mM (9/1.5 ACSF) for a robust burst rhythm or the cutting ACSF solution with K^+^ maintained at 3 mM and Ca^2+^ lowered to 1 mM (3/1 ACSF) to study burstlets and bursts together (Kam et al., 2013a). Drugs, including DAMGO, SP, picrotoxin, strychnine, and CdCl, were obtained from Sigma-Aldrich and bath-applied at the specified concentrations. In all experiments, slices were allowed to equilibrate for 30 min to ensure that the frequency and magnitude of XII and preBötC population activities reached steady-state. Respiratory activity reflecting suprathreshold action potential firing from populations of neurons was recorded from XII motor nucleus or nerve roots and as population activity directly from the preBötC using suction electrodes (tip size ~50 μm) and an Axiopatch 200A (Molecular Devices), Multiclamp 700B (Molecular Devices), and/or a differential AC amplifier (AM systems), filtered at 2–4 kHz, integrated, and digitized at 10 kHz. Integration was performed on custom built Paynter filter with a 20-100 ms time constant. Digitized data were analyzed off-line using custom procedures written for IgorPro (Wavemetrics).

### Holographic photostimulation

Holographic photostimulation was performed on a Phasor SLM system (Intelligent Imaging Innovations, Inc.) mounted around an epifluorescence upright microscope (Axioscope; Zeiss). We used a 405 nm diode laser (CUBE 405-100; Coherent) to uncage MNI-glutamate (0.5 mM) and depolarize targeted neurons. An iterative Fourier transform algorithm, implemented in Slidebook 5 (Intelligent Imaging Innovations, Inc.), computed the phase pattern on the SLM corresponding to the desired distribution of light intensity at the focal plane of the objective. A blocker was inserted in the intermediate Fourier plane to block the unmodulated light component (zero order spot) and replicate patterns (Golan et al., 2009). Neurons were targeted by centering 10 um circular spots over the somata of the selected neurons. The laser intensity per spot was set at 1-3 mW, which was previously determined to elicit bursts of action potentials in the targeted neurons without the spread of glutamate to neighboring neurons (Kam et al., 2013b). Each laser stimulation consisted of 5 × 0.8 ms pulses, delivered at 200 Hz. A threshold stimulus was determined as the minimum number of neurons and minimum laser power required to entrain preBötC bursts at slightly faster than the endogenous frequency with a success rate > 80%. The latency between laser stimulation and burst initiation was calculated as the time between the start of the next preBötC burst and the beginning of laser stimulation. preBötC activity was used here, rather than XII activity, to isolate the effects of DAMGO on rhythmogenic mechanisms and avoid additional delays or failures due to depression of XII premotoneurons and motoneurons. Successful triggering of a burst by photostimulation was defined as burst detection within 0.5 s of laser stimulation.

### Data analysis and statistics

Semiautomated event detection of respiratory-related activity recorded in XII output or preBötC population recordings was performed using custom procedures written in IgorPro. Multiple criteria, including slope and amplitude thresholds, were used to select events automatically, which were then confirmed visually. Event duration, amplitude, shape, and synchrony between XII and preBötC activity were criteria used to categorize detected events as burstlets, bursts, and doublets. Burstlets were events in preBötC that did not temporally overlap a XII burst. Doublets were distinguished from bursts by their longer duration and the presence of multiple peaks of activity. Two closely spaced bursts were considered a doublet based on the distribution of the period of XII output. A small peak at <2 s in the distribution of periods of XII bursts was usually observed. A Gaussian was fit to this small peak and a threshold time interval was set. Two bursts separated by less than this threshold were considered a doublet. As doublets were in phase with bursts and burstlets, these events were grouped with bursts and not separately analyzed.

Rhythmogenic processes were captured by measuring the frequency of preBötC rhythmic activity, which included small amplitude burstlets that were not transmitted to XII motor output, larger amplitude bursts that produced XII bursts, and longer duration doublets that were also transmitted to XII motoneurons. The average frequency was calculated as the mean of the instantaneous frequencies (1/interevent interval) across all preBötC activity (burstlets, bursts, and doublets) in that condition. Pattern generating processes that included the production and transmission of preBötC activity to XII output were characterized by measuring the fraction of preBötC events that were burstlets. The burstlet fraction was calculated as the ratio of the number of burstlets to the total number of rhythmic preBötC events (burstlets, bursts, and doublets). Amplitudes were normalized to the amplitude of the largest burst in the control condition.

Data are expressed as mean ± SD. Comparisons between two groups were performed with paired or unpaired Student’s *t*-tests. Multiple comparisons were evaluated using parametric one-way or two-way repeated measures ANOVA. Post-hoc Tukey or paired Student’s *t*-tests with the Bonferroni-Holm comparison were then used to determine significance for within group comparisons. Statistical significance was set at p < 0.05.

## Supplemental Figure Legends

**Figure S1. μOR, TAC1, and NK1R expression in preBötC.** Related to Figure 3.

(A) Confocal image of a medullary brainstem section at the level of the preBötC from a P2 wild type mouse triple processed with RNAScope. Probes for NK1R (green), μOR (magenta), and TAC1 (white) were used, and the tissue was counterstained with DAPI (blue). Scale bar, 20 μm.

(B) Higher magnification images showing colocalization of TAC1, NK1R, and μOR puncta in several preBötC neurons (arrows) as well as a neuron that expressed NK1R and μOR, but not TAC1 (arrowhead). Scale bar, 25 μm.

**Figure S2. Substance P increases preBötC *f* in *Dbx1*^*Cre*^;*Oprm1*^*fl/fl*^ mice.** Related to Figure 4.

(A) Representative traces showing ∫preBötC and ∫XII population activity from a *Dbx1*^*Cre*^;*Oprm1*^*fl/fl*^ slice in control (ctrl, left), 100 nM DAMGO (middle), and 500 nM Substance P (SP, right). Scale bar, 10 s.

(B) SP (500 nM) significantly increased preBötC *f* on slices from *Dbx1*^*Cre*^;*Oprm1*^*fl/fl*^ mice, while 100 nm DAMGO had no effect *, p < 0.05, One-way ANOVA post-hoc Tukey test, n=4.

