## Supplemental Figures for "Inspiratory rhythmogenic activity is burst-independent and opioid-sensitive"

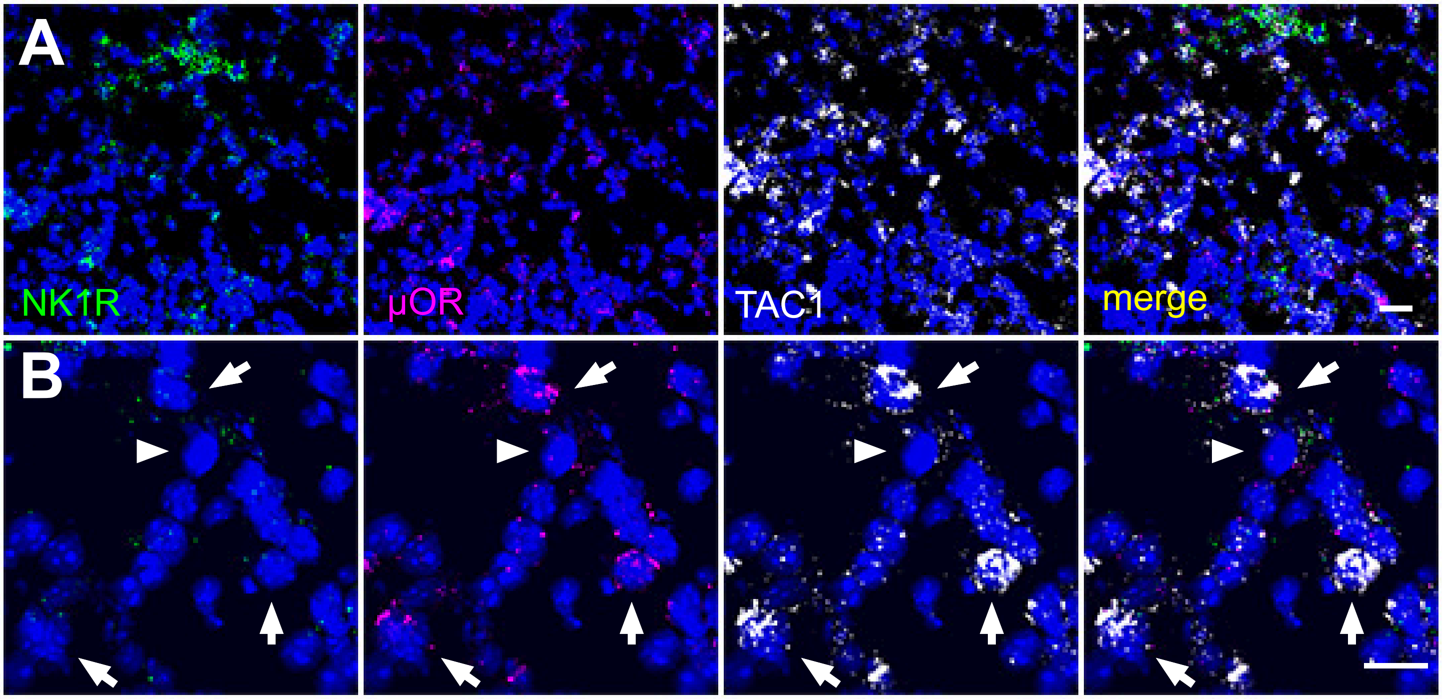


**Figure S1. µOR, TAC1, and NK1R expression in preBötC.** Related to Figure 3.
(A) Confocal image of a medullary brainstem section at the level of the preBötC from a P2 wild type mouse triple processed with RNAScope. Probes for NK1R (green), µOR (magenta), and TAC1 (white) were used, and the tissue was counterstained with DAPI (blue). Scale bar, 20 µm.
(B) Higher magnification images showing colocalization of TAC1, NK1R, and µOR puncta in several preBötC neurons (arrows) as well as a neuron that expressed NK1R and µOR, but not TAC1 (arrowhead). Scale bar, 25 µm.

Figure S1

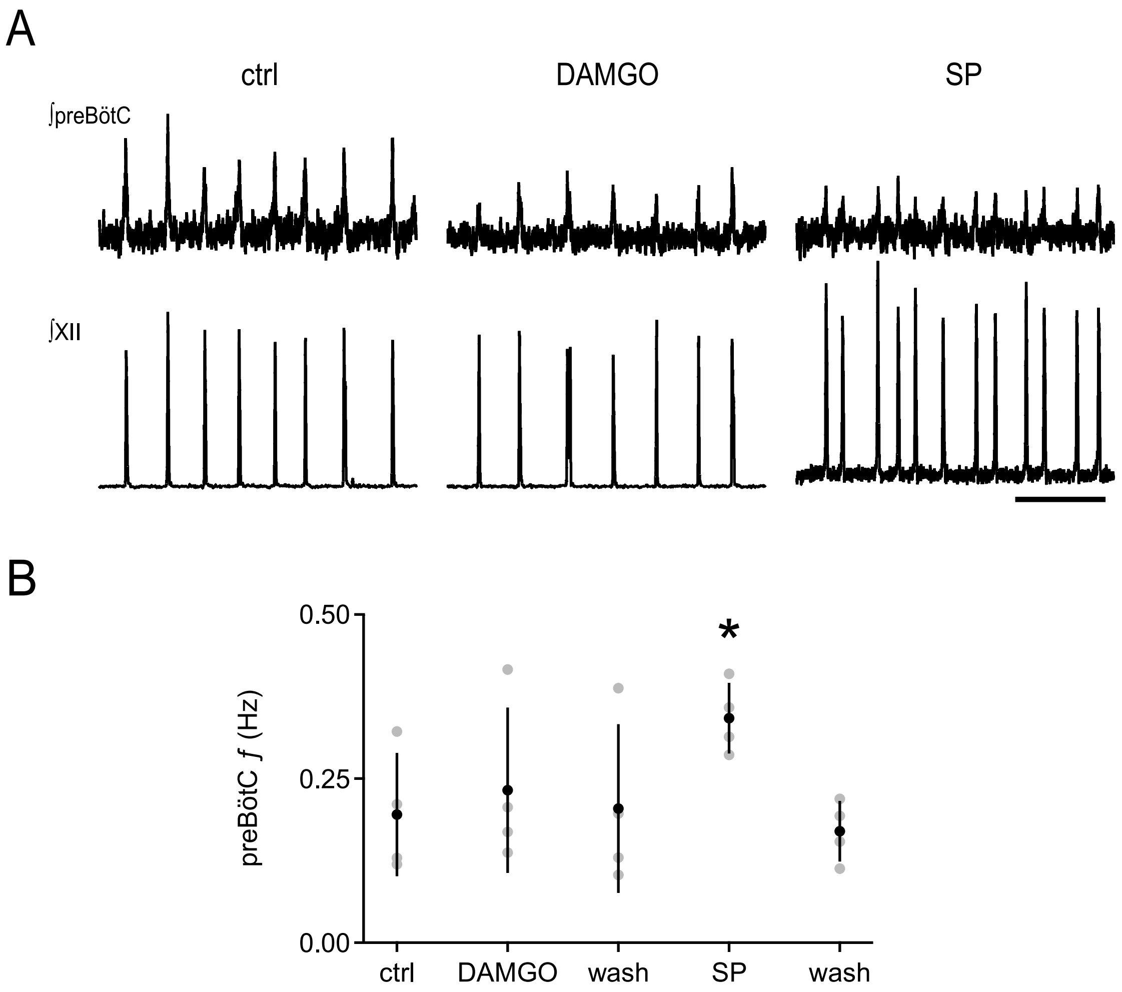


**Figure S2. Substance P increases preBötC ƒ in *$Dbx1^{cre}$;$Oprm1^{fl\backslash fl}$ mice.*** Related to Figure 4. (A) Representative traces showing ∫preBötC and ∫XII activity from a *$Dbx1^{cre}$;$Oprm1^{fl\backslash fl}$* slice in control (ctrl, left), 100 nM DAMGO (middle), and 500 nM Substance P (SP, right). Scale bar, 10 s. (B) SP (500 nM) significantly increased preBötC ƒ in slices from *$Dbx1^{cre}$;$Oprm1^{fl\backslash fl}$* mice while 100 nM DAMGO had no effect. *, $p<0.05$, One-way ANOVA, post-hoc Tukey test, n=4.

Figure S2
